# PSMAx-Guided PROTAC Degraders for Tumor-Specific Protein Degradation in Prostate Cancer

**DOI:** 10.1101/2024.04.25.591100

**Authors:** Xiaolei Meng, Xiaolin Hu, Siqi Zhang, Sai Zhang, Xiao Wang, Shumin Ma, Chong Qin

## Abstract

PROTACs, degrading target protein to treat diseases, represent a highly promising drug design strategy. However, the degradation of target proteins by PROTACs in non-disease tissues may lead to systemic toxicity. Herein, capitalizing on the characteristic overexpression of PSMA in prostate cancer tumor tissues, we devised a PSMA-guided PROTACs specific targeting to prostate cancer. By conjugating AR degraders and BET degraders separately with PSMA ligands via cleavable linkers, two classes of PSMA-guided PROTACs were obtained. *In vitro* experiments demonstrated that PSMA-guided PROTAC molecules selectively degraded target proteins in PSMA-overexpressing prostate cancer cells, without affecting target proteins in non-PSMA-overexpressing cells. *In vivo* studies revealed that compared to conventional PROTACs, PSMA-guided PROTACs enhanced drug exposure in prostate cancer tumor tissues, prolonged half-life, and consequently achieved stronger and more sustained therapeutic effects. The PSMA-guided PROTAC strategy provides a novel avenue for disease tissue-specific PROTAC research, holding significant implications for targeted therapy in prostate cancer.

## Introduction

Prostate cancer (PC) is the most prevalent malignancy among men and the second leading cause of cancer-related death.^1–3^ Conventional treatment, such as surgery, radiation, and androgen suppression are effective in early-stage localized prostate cancer.^4,5^ Unfortunately, a considerable proportion of patients treated with androgen deprivation therapy recurrence of androgen-independent cancer, progressing into castration-resistant prostate cancer (CRPC).^6–8^ There is an urgent imperative for the development of specific and targeted agents for the treatment of CRPC.

PROTACs are bifunctional molecules comprised of three components: the first being a ligand that binds to the target protein, the second being a ligand that binds to the E3 ligase, with a linker in between.^9^ Their mechanism of action involves recruiting the target protein and E3 ligase together to form a ternary complex, whereby ubiquitin is then transferred onto the target protein.^10–12^ Subsequently, the proteasome degrades the target protein by recognizing the ubiquitin moiety on it.^13^ As a novel approach, PROTACs has demonstrated promising and notable advantages in the treatment of CRPC.^14–20^ **ARV-110**^21^ and **ARV-766**^22^ are two orally bioavailable androgen receptor (AR) degraders currently undergoing clinical trial investigations for the treatment of CRPC. Recently, we have reported **BWA-522**, a degrader targeting the N-terminal domain of AR, capable of simultaneously degrading both full-length AR and AR splice variants.^20^ In addition to AR degraders, the Crew’s group developed **ARV-771** (Figure 1A), a bromodomain and extraterminal domain (BET) degrader. **ARV-771** could effectively suppress the BET protein levels and AR-mediated gene transcription, leading to significant tumor regression in a xenograft model of CRPC. ^23^

**Figure 1.**
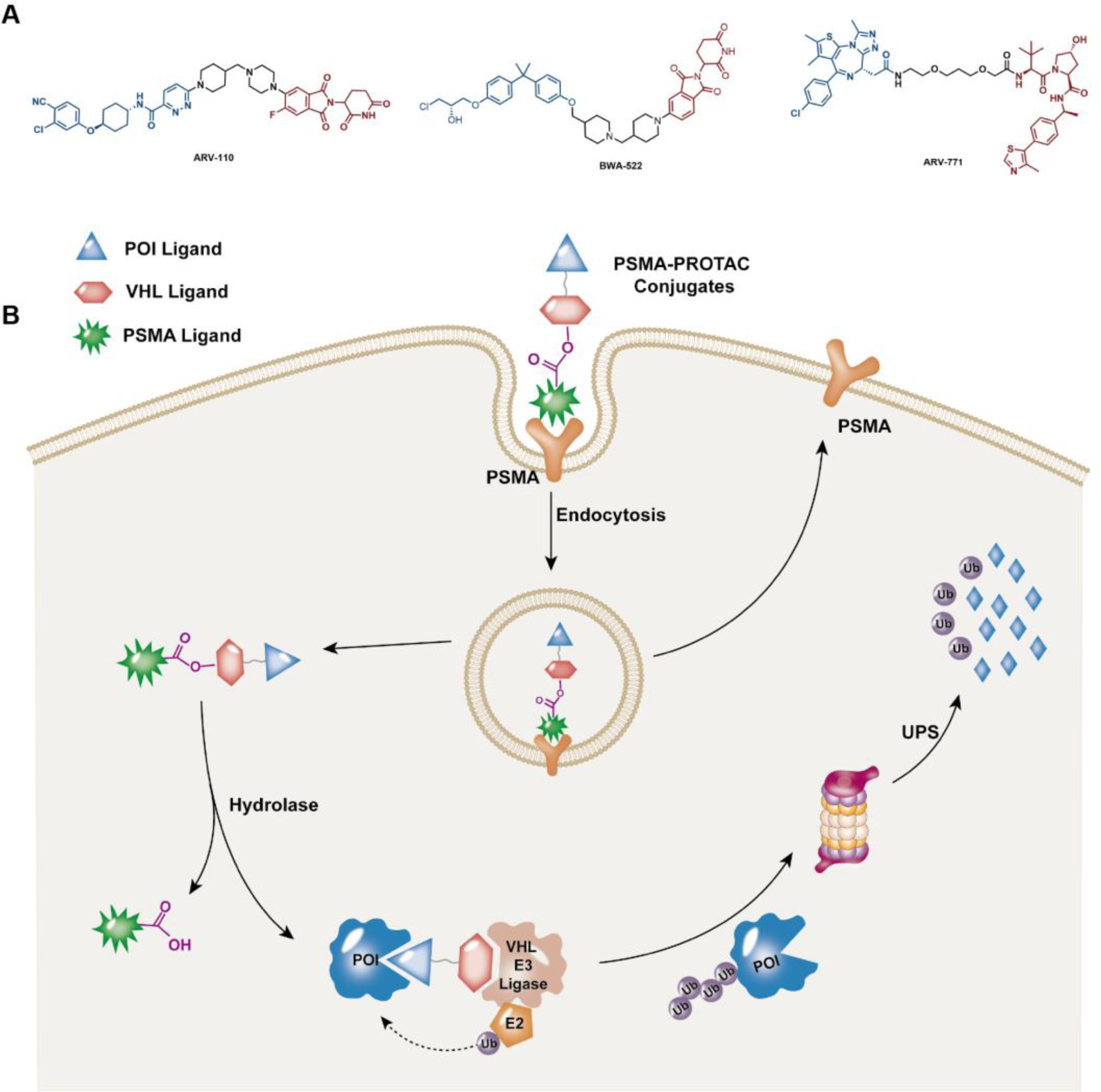
PSMA transports PSMA-PROTAC conjugates into prostate cancer cells for the selective degradation of POIs. (A) PROTACs in clinical trials or research for the treatment of CRPC. (B) Schematic representation of the PSMA-PROTAC strategy. After the PSMA ligand of PSMA-PROTAC conjugate recognizes the PSMA receptor on the surface of prostate cancer cells, the conjugates are rapidly internalized into prostate cancer cells where they release PROTACs in the presence of intracellular hydrolases, leading to the degradation of POIs in the ubiquitin–proteasome pathway.

While PROTAC degraders have made significant progress in the treatment of CRPC, degradation of target proteins outside prostate cancer tumor tissues may lead to potential off-tissue effect and systemic toxicity, thereby reducing the therapeutic window of the drugs.^24,25^ Therefore, the discovery of degraders with the ability to specifically target prostate cancer tissues is of paramount importance. Prostate-specific membrane antigen (PSMA) is a transmembrane zinc metalloprotease classified as a type II transmembrane protein, also known as glutamate carboxypeptidase II.^26,27^ The expression of PSMA is significantly upregulated in prostate cancer, whereas its levels remain comparatively lower in healthy prostate and other tissues.^28,29^ Notably, a positive correlation has been observed between PSMA levels and prostate tumor aggressiveness.^30^ As such, PSMA has emerged as a promising target for prostate cancer drug delivery, diagnostics, and intraoperative navigation.^31–36^ In this study, we designed and synthesized PSMA-guided PROTAC degraders, which are preferentially transported into PSMA-overexpressing prostate cancer cells (Figure 1B). PSMA-guided degraders were generated by tethering VHL-based degraders and PSMA ligands with an esterolysis-cleavable linker. These conjugates could be internalized into prostate cancer cells by PSMA^37^ and then hydrolyzed by intracellular hydrolases^38–40^ and release active degraders. This strategy facilitates the targeted delivery of PROTACs to PSMA-overexpressing prostate cancer cells, thereby enabling the selective protein degradation in prostate cancer tissue while minimizing off-target effects.

## Results and Discussion

### Design and Synthesis of the PSMA-guided AR Degraders

As the primary relevant protein in prostate cancer, we selected the androgen receptor (AR) as the initial target for our research.^41^ We had successfully discovered a VHL-based AR degrader, **ARD-203** (Figure 2), which exhibited exceptional degradation activity *in vitro*. The hydroxyl group located on the hydroxyproline group plays a crucial role in recruiting the Cul2-VHL E3 ubiquitin ligase complex^42^, and chemical modifications to this hydroxyl group lead to a loss of activity. A widely used PSMA ligand (glutamate-urea-lysine) was tethered to the hydroxyl group via an ester bond (Figure 2), generating **PSMA-ARD-203**. Before entering prostate cancer cells, the **PSMA-ARD-203** remains in an inert, non-activated form. After specific internalization into prostate cancer cells, active degrader **ARD-203** was released in the presence of intracellular hydrolases.

**Figure 2.**
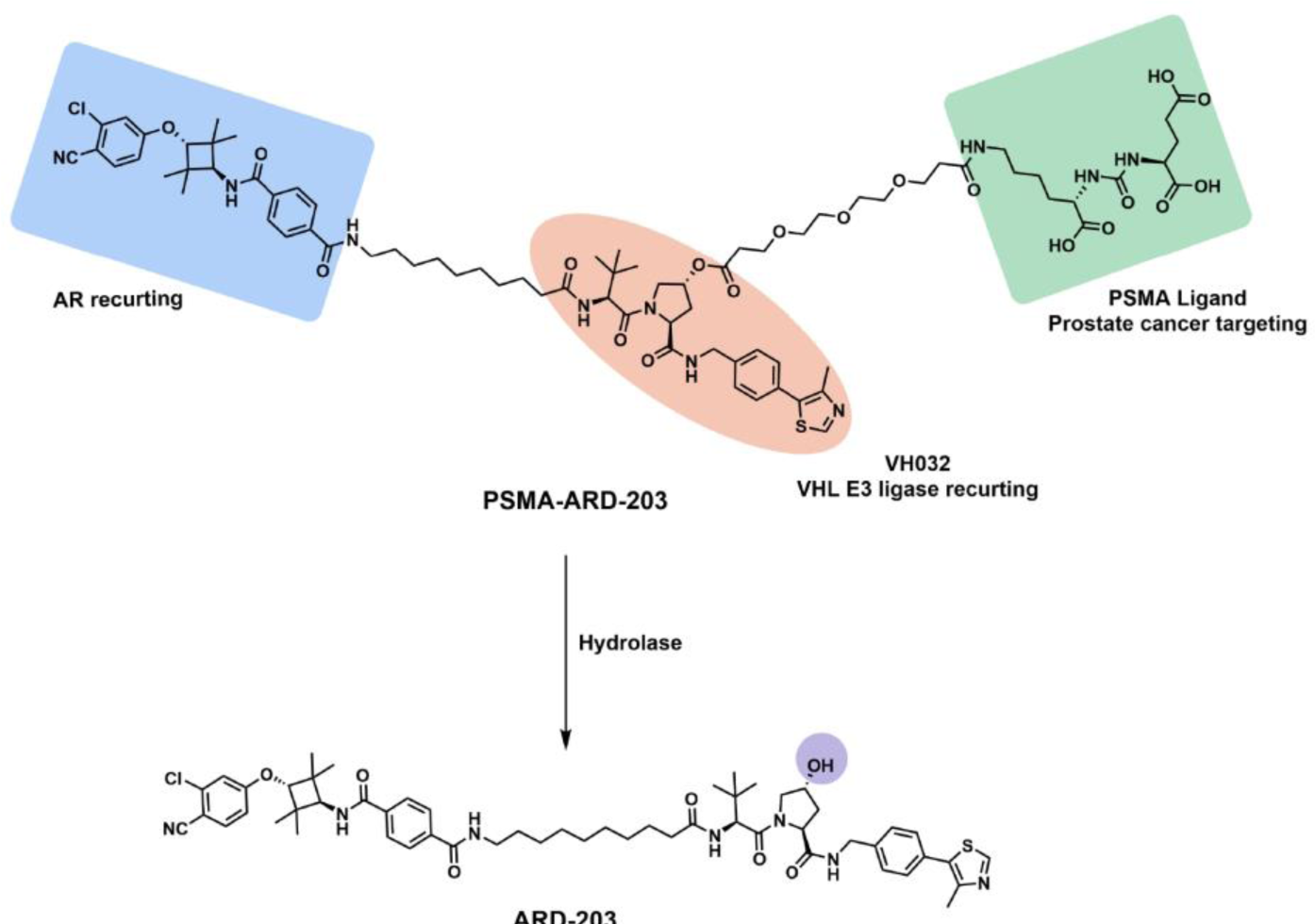
Design of **PSMA-ARD-203**. Schematic illustration of the activation of **PSMA-ARD-203** by intracellular hydrolase.

We tested the stability of **PSMA-ARD-203** in various media, including phosphate buffered saline (PBS), Dulbecco’s Modified Eagle Medium (DMEM) basal media, or DMEM supplemented with 10% fetal bovine serum at 37°C. The results indicated that **PSMA-ARD-203** exhibited stability over 90% for up to 24 h at the tested conditions (Figure S1).

### PSMA-ARD-203 Specifically Degrades AR in Prostate Cancer Cells

To determine the specific role of **PSMA-ARD-203** in prostate cancer cells versus that in non-prostate cancer cells, three prostate cancer cell lines with high PSMA expression, namely LNCaP cells, 22Rv1 cells, and VCaP cells, as well as three non-prostate cell lines with low PSMA expression, namely human breast carcinoma cells (MCF-7 cells and MDA-MB-231 cells) and human embryonic kidney cells (HEK-293T cells) were chosen (Figure S2A). Western blotting of the cytosolic and membrane fractions from 22Rv1 cells and HEK-293T cells indicated that PSMA was mainly expressed on the membrane in 22Rv1 cells but was not detected in HEK-293T cells (Figure S2B).

To investigate the AR-degradation activity of **PSMA-ARD-203**, western blotting was conducted with different concentrations of **PSMA-ARD-203**, with **ARD-203** as a control. The efficiency of **PSMA-ARD-203** in inducing AR degradation in three prostate cancer cell lines with high PSMA expression was comparable with that of **ARD-203** (Figure 3A). **PSMA-ARD-203 a**efficiently degraded AR in a concentration-dependent manner in VCaP, LNCaP and 22Rv1 cells (Figure 3B), and the DC_50_ values for these cell lines were determined to be 21.86 ± 13.40 nM, 44.38 ± 18.05 nM, and 50.19 ± 13.78 nM, respectively (Figure 3C). In comparison, **PSMA-ARD-203** showed only slight influences on AR protein levels in MCF-7, MDA-MB-231 and HEK-293T cells (Figure 3B), with DC_50_ values exceeding 5 μM (Figure S3). These results indicated that **PSMA-ARD-203** exhibited remarkable efficiency in specifically degrading AR in PSMA-overexpressing prostate cancer cells compared with that in non–prostate cancer cells with low PSMA expression.

**Figure 3.**
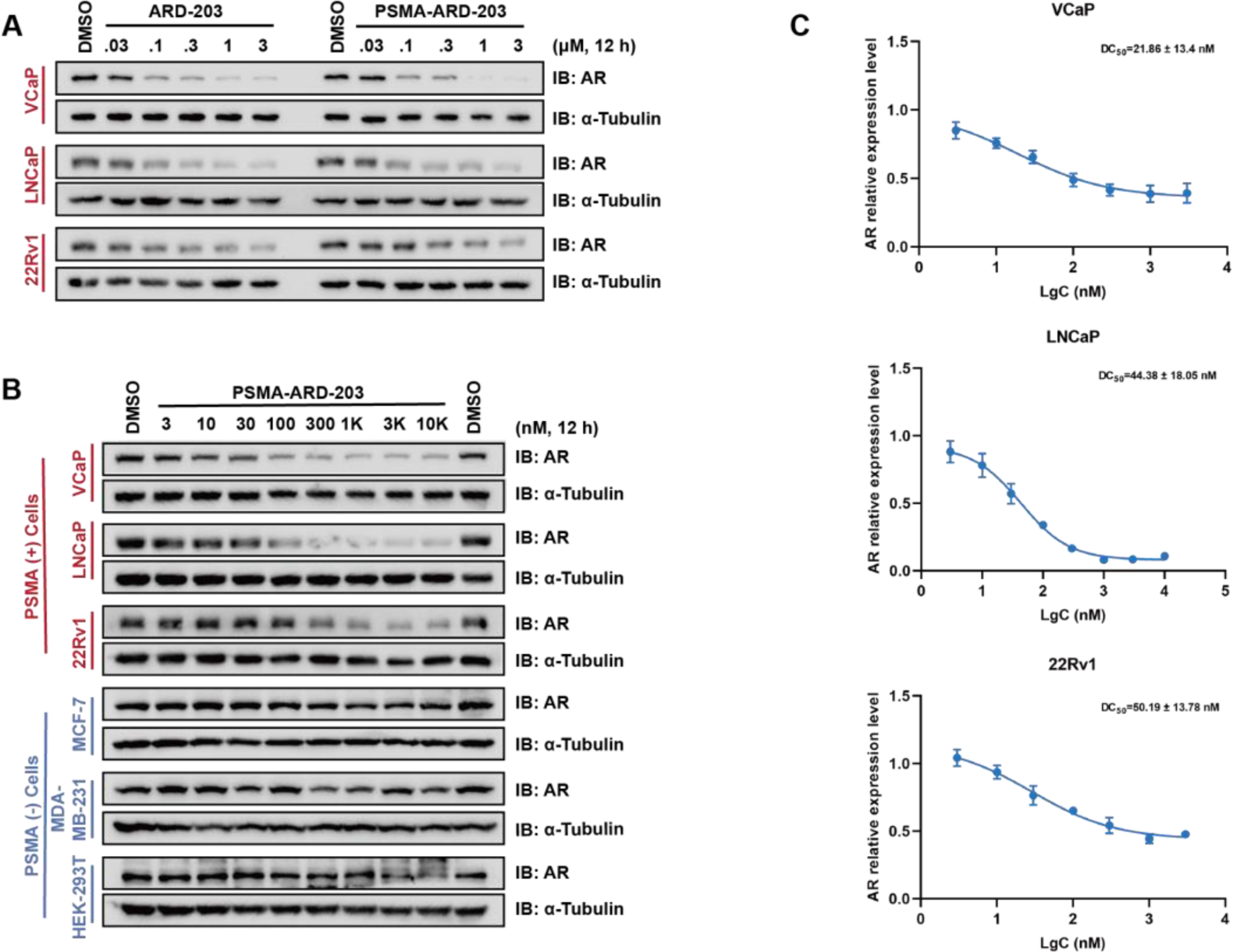
**PSMA-ARD-203** preferentially degrades AR in prostate cancer cells. (A) Western blotting of AR levels in VCaP, LNCaP and 22Rv1 cells treated with the indicated concentrations of **ARD-203** or **PSMA-ARD-203** for 12 h. (B) Western blotting of AR levels in prostate cancer cells or non–prostate cancer cells treated with the indicated concentrations of **PSMA-ARD-203** for 12 h. (C) DC_50_ values of **PSMA-ARD-203** based on quantification of AR levels normalized to α-Tubulin levels or GAPDH levels in prostate cancer cell lines in B (n=3). Data are presented as mean ± standard error of the mean (n=3).

### PSMA-ARD-203 Degrades AR in a VHL- and Proteasome-dependent Manner

To validate that **PSMA-ARD-20**3–induced degradation of AR depends on VHL E3 ubiquitin ligase, CRPC cells were co-treated with the VHL ligand VH032 and **PSMA-ARD-203**. Co-treatment with VH032 effectively inhibited AR-degradation activity in LNCaP and 22Rv1 cells (Figure 4A, 4B). Furthermore, the proteasome inhibitor MG132 and Cullin-RING ubiquitin ligase (CRL) E3 ligase inhibitor MLN4924 also blocked the effect of **PSMA-ARD-203** in degrading AR in LNCaP and 22Rv1 cells (Figure 4C, 4D). Collectively, these results demonstrated that the PSMA-guided AR degrader, **PSMA-ARD-203**, degraded AR in prostate cancer cells in a VHL- and proteasome-dependent manner.

**Figure 4.**
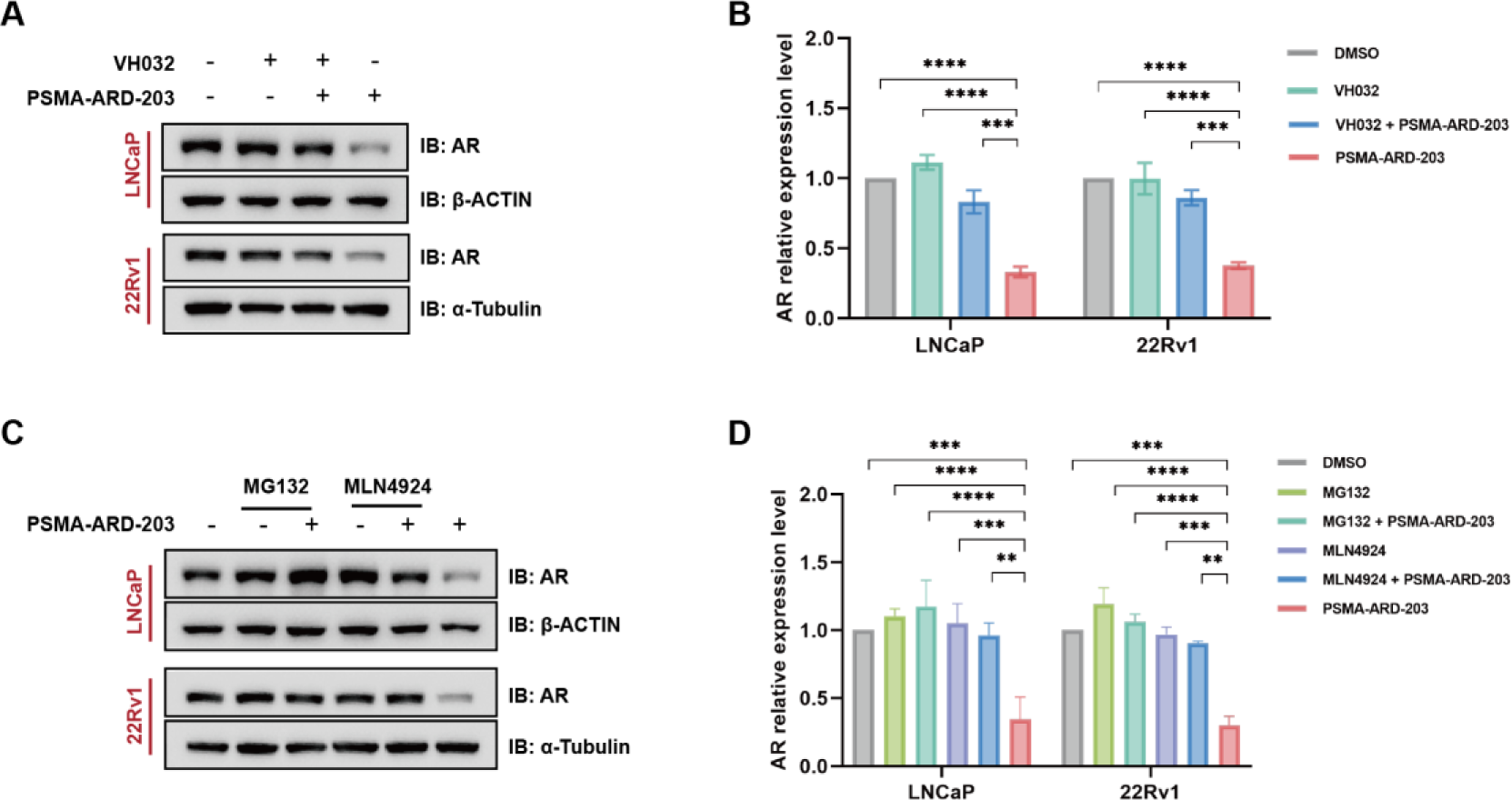
**PSMA-ARD-203** degrades AR in a VHL- and proteasome-dependent manner. (A) Western blotting of AR levels in LNCaP and 22Rv1 cells pretreated with 200 μM VH032 for 1 h followed by treatment with 1 μM **PSMA-ARD-203** for 6 h. (B) Quantification of AR levels from the western blots in A (n=3). (C) Western blotting of AR levels in LNCaP and 22Rv1 cells after treatment with 4 μM MG132 or 1 μM MLN4924 and 1 μM **PSMA-ARD-203** for 8 h. (D) Quantification of AR levels using the western blots in C (n=3). Data are presented as mean ± standard error of the mean (n=3). \**p*<0.05, \*\**p*<0.01, \*\*\**p*<0.001, \*\*\*\**p*<0.0001, two-way analysis of variance.

### PSMA-ARD-203 Degrades AR in a PSMA-dependent Manner

To validate the significance of PSMA in facilitating the targeted delivery of **PSMA-ARD-203**, LNCaP and 22Rv1 cells were pretreated with the PSMA ligand. As anticipated, the AR-degradation activity of **PSMA-ARD-203** was blocked by the PSMA ligand (Figure 5A, S4A). Furthermore, PSMA knockdown by siRNAs attenuated **PSMA-ARD-203**–induced AR degradation in LNCaP cells (Figure 5B, S4B).

**Figure 5.**
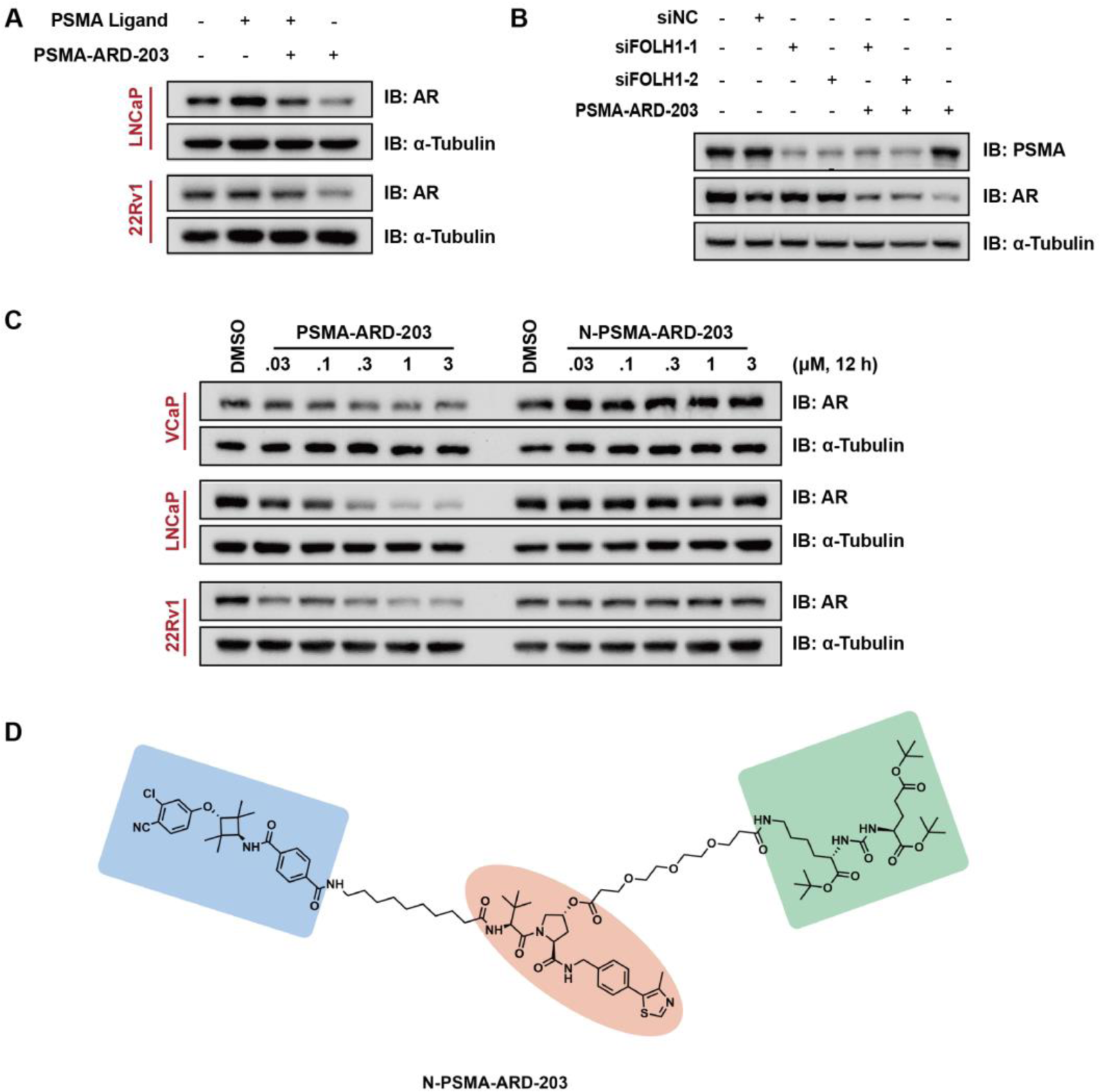
**PSMA-ARD-203** degrades AR in a PSMA-dependent manner. (A) Western blotting of AR levels in LNCaP and 22Rv1 cells pretreated with 200 μM PSMA ligand for 2 h followed by treatment with 1 μM **PSMA-ARD-203** for 6 h. (B) LNCaP cells with or without knockdown of endogenous PSMA were treated with 1 μM **PSMA-ARD-203** for 24 h and AR levels were determined using western blotting. (C) Western blotting of AR levels in VCaP, LNCaP, and 22Rv1 cells treated with indicated concentrations of **PSMA-ARD-203** or **N-PSMA-ARD-203** for 12 h. (D) Structure of the negative control (**N-PSMA-ARD-203**) of **PSMA-ARD-203**.

To obtain a negative control compound, the three carboxyl groups in the PSMA ligand were protected with tert-butyl groups yielding **N-PSMA-ARD-203** (Figure 5D). This compound has lost its ability to bind with PSMA, thus preventing internalization into cells through PSMA. As expected, **N-PSMA-ARD-203** was ineffective in degrading AR in the three prostate cancer cell lines (Figure 5C, S4C, S5). These results indicated that **PSMA-ARD-203** entered prostate cancer cells and degraded AR in a PSMA-dependent manner.

### PSMA-ARD-203 Accumulates within 22Rv1 Xenograft Tumors *in vivo*

To assess whether **PSMA-ARD-203** exhibited *in vivo* tumor-targeting ability, its pharmacokinetics (PK) in 22Rv1 xenograft tumor–bearing mice was determined after intravenous administration (0.3 mmol/L) and using **ARD-203** as a control (Figure 6A). As shown in Figure 6B, **PSMA-ARD-203** showed higher drug concentrations in tumors versus **ARD-203** at all the time points. The effective and gradual release of **ARD-203** by **PSMA-ARD-203** was also observed. Furthermore, a higher concentration of PSMA-**ARD-203** and its **released-ARD-203** were still detected in tumors 96 h post-injection, whereas no drug was noted in the tumors of **ARD-203**–treated mice. Compared with **ARD-203**, **released-ARD-203** had higher *in vivo* exposure to the tumor and plasma (Figure 6C, Table S1). These results demonstrated the exceptional tumor-targeting capabilities of **PSMA-ARD-203** in the 22Rv1 xenograft model.

**Figure 6.**
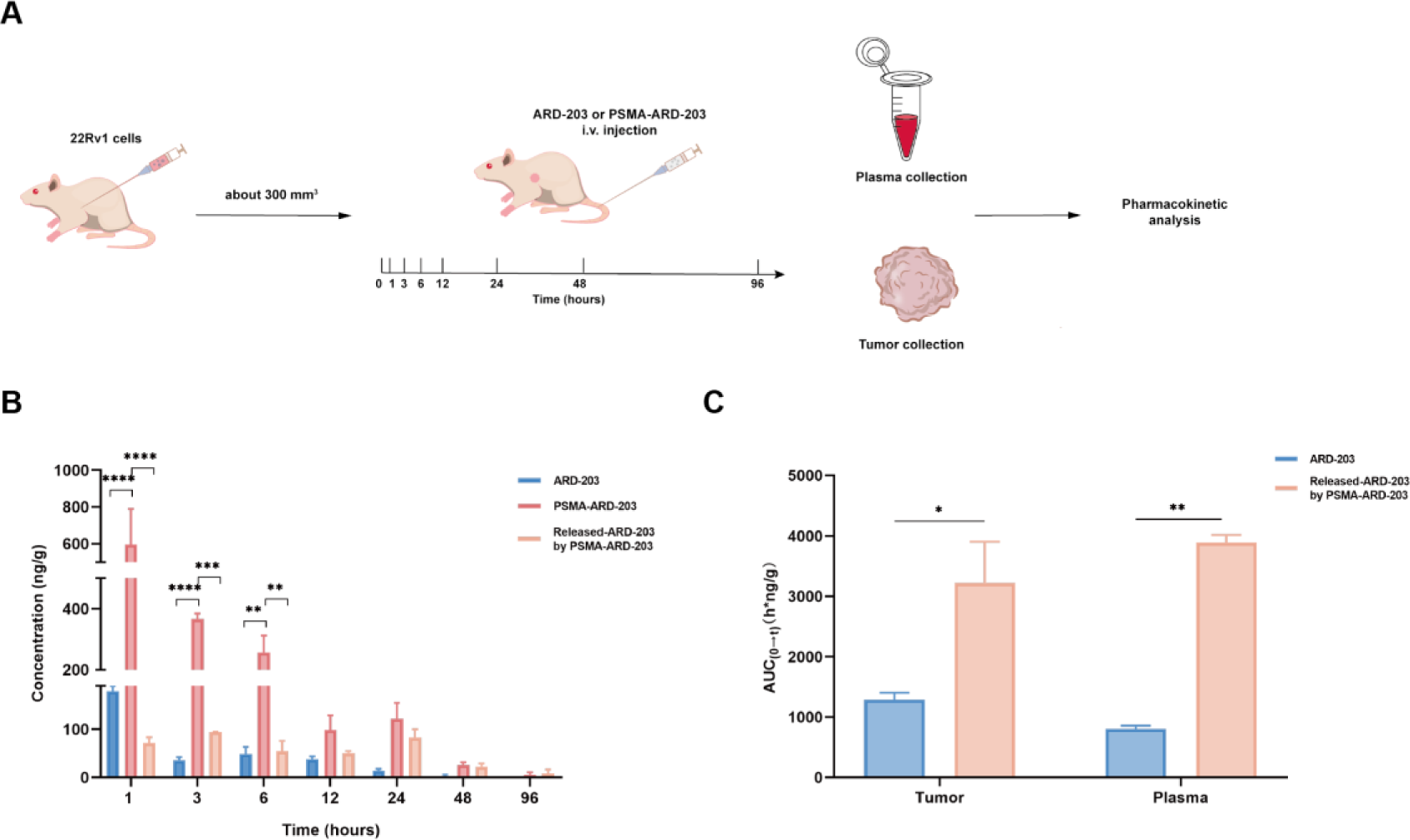
Ability of **PSMA-ARD-203** and **ARD-203** to enter 22Rv1 xenograft tumors. (A) Schematic depiction of pharmacokinetic analysis in 22Rv1 xenograft tumor-bearing mice after the treatment **ARD-203** or **PSMA-ARD-203** with intravenous administration at 0.3 mmol/L. (B) Drug concentration–time profiles of **ARD-203**, **PSMA-ARD-203**, and its hydrolysis product in 22Rv1 xenograft tumors (n=3). (C) Comparison of the area under the curve (AUC)_0-t_ of **ARD-203** and released-**ARD-203** from **PSMA-ARD-203** in the tumor and plasma (n=3). Data are presented as mean ± standard error of the mean (n=3). \**p*<0.05, \*\**p*<0.01, \*\*\**p*<0.001, \*\*\*\**p*<0.0001, two-way analysis of variance.

The more comprehensive PK data are summarized in Table S1. Intravenous administration **PSMA-ARD-203** led to a high exposure of 7112.00 ± 1572.00 h·ng/g, maximum concentration of 597.00 ± 332.00 ng/g, low clearance of 628.00 ± 133.80 g/kg/h. The PK profile of **released-ARD-203** from **PSMA-ARD-203** in tumors was determined for a more direct comparison with **ARD-203**. **Released-ARD-203** had an excellent overall PK profile with a higher area under the curve (AUC)_0-t_ (3223.00 ± 1182.00 h·ng/g), lower clearance (1330.00 ± 652.80 g/kg/h), reasonably longer half-life (T_1/2_) of 24.80 ± 17.63 h and MRT_0-t_ of 24.18 ± 8.35 h. These results indicated that **PSMA-ARD-203** could enhance drug distribution in tumors, increase drug exposure, reduce the drug clearance rate from tumors, facilitate the slow release of drugs in tumors, and prolong the duration of drug action.

### PSMA-ARD-203 Effectively Degrades AR in LNCaP Xenograft Tumors *in vivo*

Based on the promising PK profile in tumors, pharmacodynamic (PD) studies of **PSMA-ARD-203** were conducted to evaluate its efficacy in reducing AR protein levels in LNCaP xenograft tumors in mice and by using **ARD-203** as the control (Figure 7A, 7B). Western blotting showed that the intravenous administration of 1 mmol/L **PSMA-ARD-203** for 3 days effectively induced AR degradation in LNCaP tumors. Notably, **PSMA-ARD-203** was significantly more effective in reducing AR protein levels in LNCaP tumors compared with **ARD-203** at both the 48 h and 72 h time points. These results demonstrated that **PSMA-ARD-203** could accumulate in LNCaP tumor tissues, subsequently releasing **ARD-203** in the presence of intracellular hydrolase, which induced AR degradation in tumor tissues.

**Figure 7.**
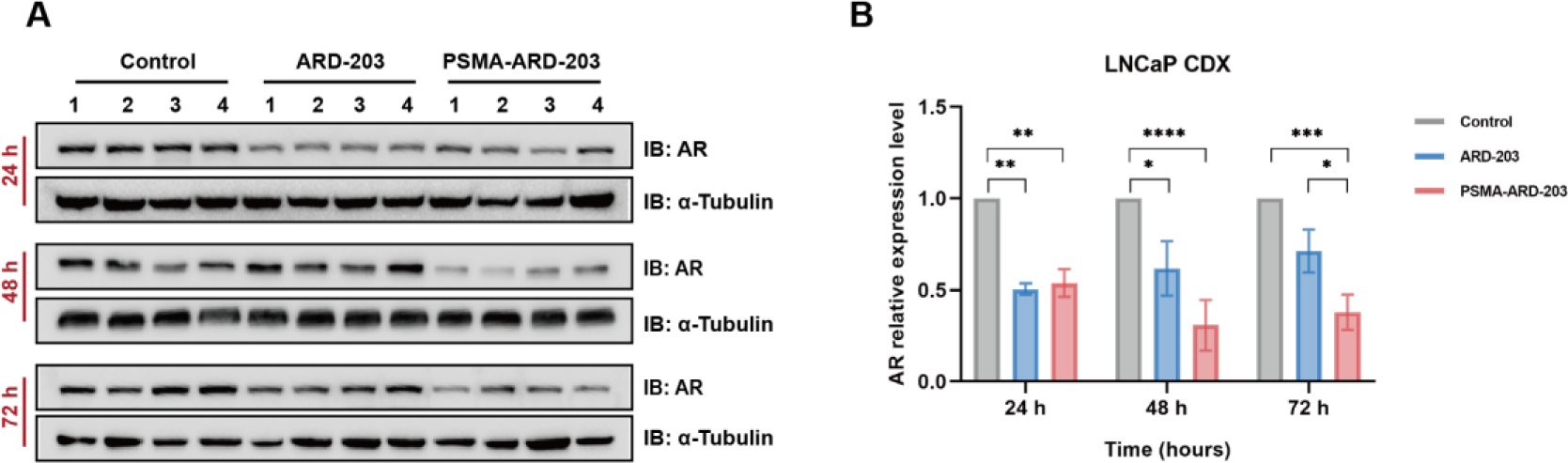
Pharmacodynamic studies of **ARD-203** or **PSMA-ARD-203** in the LNCaP xenograft model of mice. LNCaP tumor–bearing mice were treated with 1 mmol/L **PSMA-ARD-203** or **ARD-203** intravenously for 3 days. Mice were euthanized at the indicated time points for the collection of tumor tissues for analysis. (A) Western blotting of AR proteins in LNCaP tumors. (B) Quantification of AR proteins from the western blots in A (n=4). Data are presented as mean ± standard error of the mean (n=4). \**p*<0.05, \*\**p*<0.01, \*\*\**p*<0.001, \*\*\*\**p*<0.0001, two-way analysis of variance.

### PSMA-ARV-771 Specifically Degrades BET Protein in Prostate Cancer Cells

To further investigate the suitability of PSMA-guided strategy to other VHL-based PROTACs, we designed and synthesized a PSMA-guided BET protein degrader (**PSMA-ARV-771**), based on a well-studied BET protein degrader, **ARV-771**^23^ (Figure 8). The design of this conjugate involved a novel PSMA ligand with significantly enhanced affinity (IC_50_=9 ± 3 nM).^43–46^ Similar to **PSMA-ARD-203**, **PSMA-ARV-771** could be cleaved by intracellular hydrolase to release **ARV-771**, degrading BET proteins.

**Figure 8.**
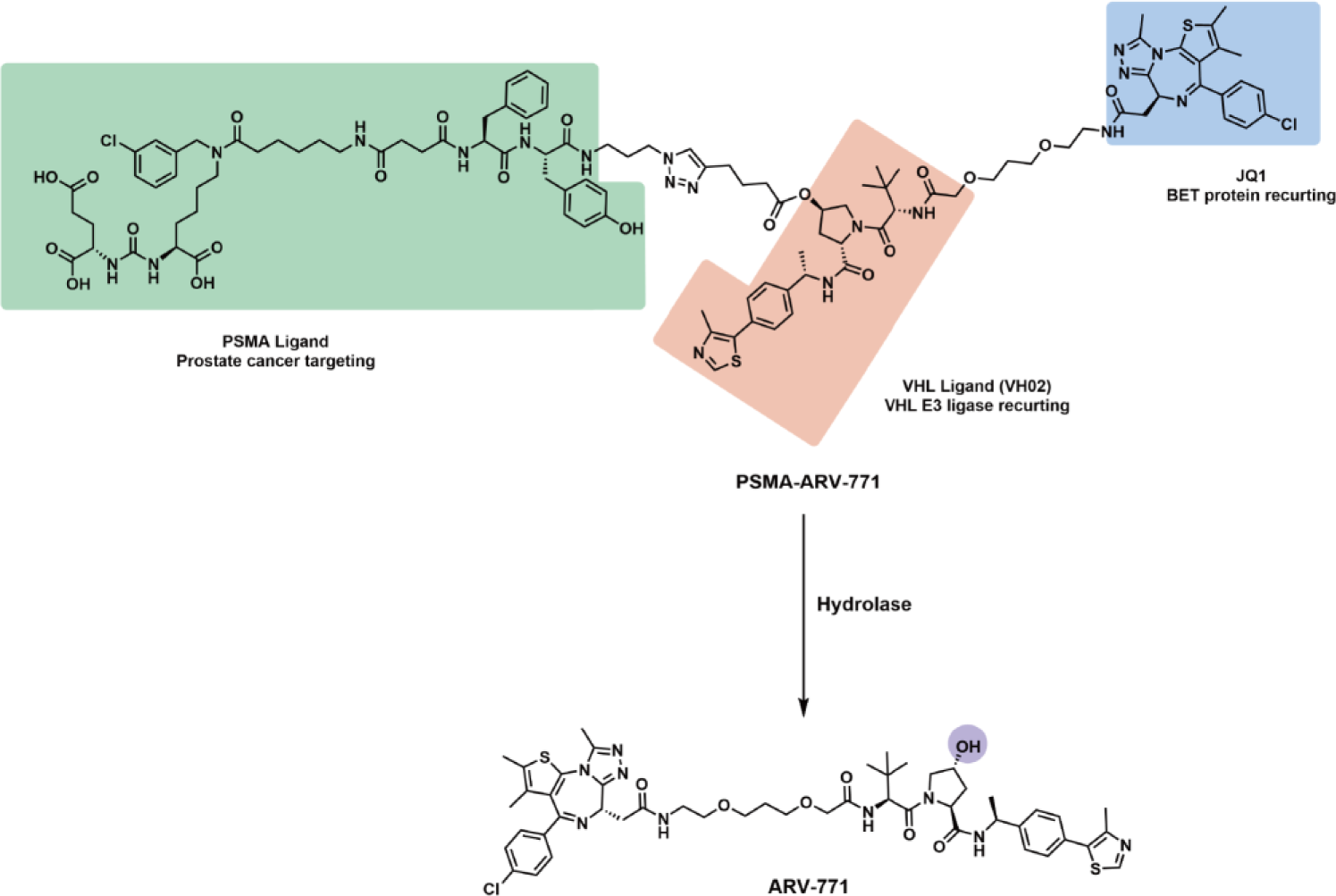
Design of **PSMA-ARV-771**. Schematic of the activation of **PSMA-ARV-771** by intracellular hydrolase.

We assessed the efficiency of **PSMA-ARV-771** in degrading BRD4 protein in PSMA-overexpressing prostate cancer cells, and cancer cells or normal cells with low PSMA expression, using **ARV-771** as a control. As seen in Figure 9A, **PSMA-ARV-771** degraded the BRD4 protein in a dose-dependent manner in VCaP, LNCaP and 22Rv1 cells as efficiently as **ARV-771**. Consistent with these findings, **PSMA-ARV-771** demonstrated a cell-killing IC_50_ comparable with that of **ARV-771** in three prostate cancer cell lines (Figure S6). In contrast, the efficacy of **PSMA-ARV-771** in degrading the BRD4 protein was relatively lower in HEK-293T, DU145 and J82 cells. The knockdown of endogenous PSMA also abrogated the effect of **PSMA-ARV-771** on BRD4 degradation in LNCaP cells (Figure 9B, 9C). Furthermore, co-treatment with VHL ligand (VH032), the proteasome inhibitor MG132 and Cullin-RING ubiquitin ligase (CRL) E3 ligase inhibitor MLN4924 blocked the degradation of BRD4 protein in LNCaP cells (Figure 9D, 9E), indicating that **PSMA-ARV-771** degraded BRD4 protein in a VHL- and proteasome-dependent manner. Collectively, these findings demonstrated that **PSMA-ARV-771** specifically degraded the BRD4 protein in prostate cancer cells having high PSMA expression.

**Figure 9.**
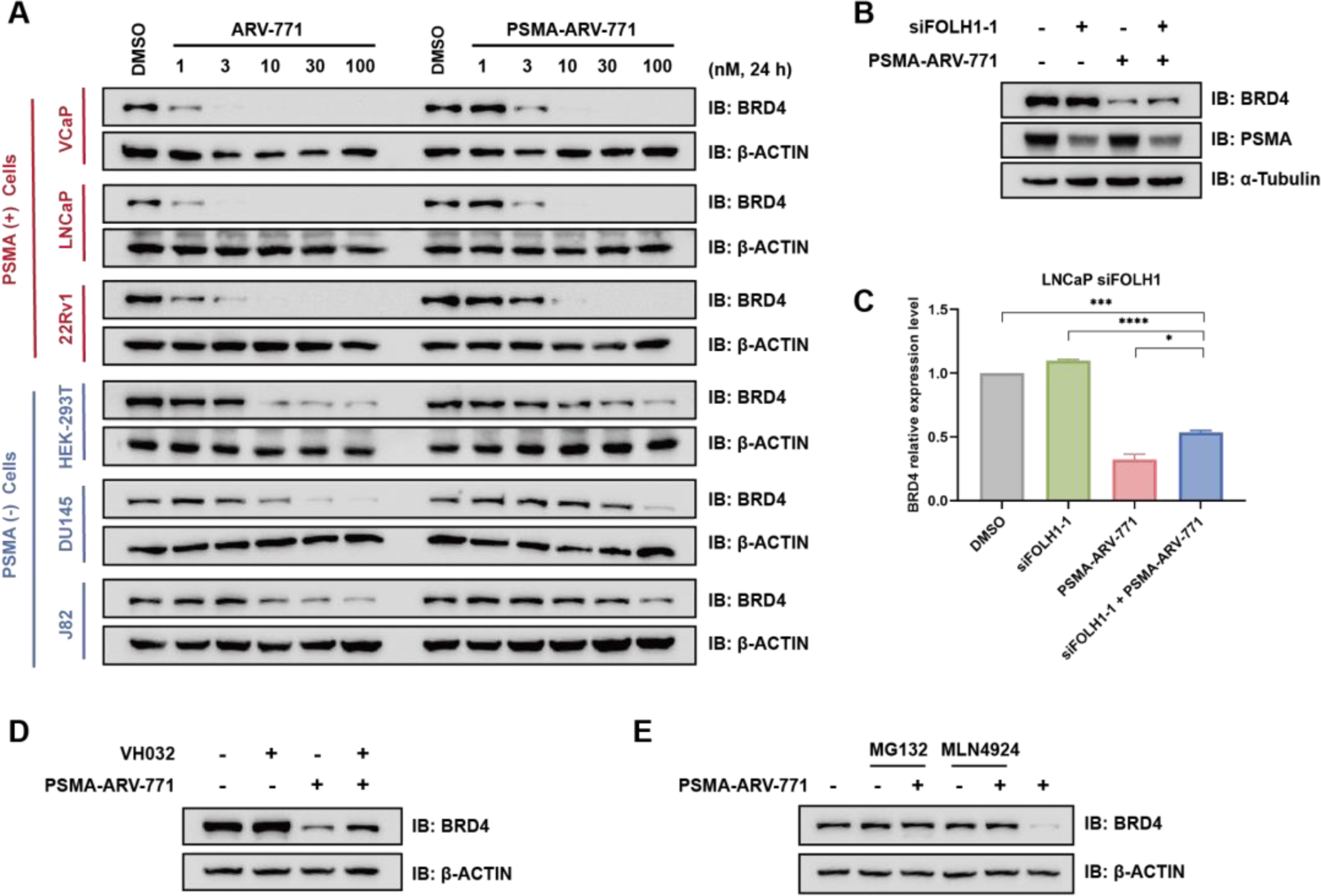
**PSMA-ARV-771** specifically degrades BRD4 protein in prostate cancer cells. (A) Western blotting of BRD4 levels in PSMA-overexpressing prostate cancer cells, and cancer cells or normal cells with low PSMA expression treated with the indicated concentrations of **ARV-771** or **PSMA-ARV-771** for 24 h. (B) LNCaP cells with or without knockdown of endogenous PSMA were treated with 3 nM **PSMA-ARV-771** for 12 h, and BRD4 levels were determined using western blotting. (C) Quantification of BRD4 levels using the western blots in B (n=3). (D) Western blotting of BRD4 levels in LNCaP cells pretreated with 100 μM VH032 for 1 h followed by co-treatment with 10 nM **PSMA-ARV-771** for 8 h. (E) Western blotting of BRD4 levels in LNCaP cells pretreated with with 4 μM MG132 or 1 μM MLN4924 for 1 h followed by co-treatment with 10 nM **PSMA-ARV-771** for 8 h. Data are presented as mean ± standard error of the mean (n=3). \**p*<0.05, \*\**p*<0.01, \*\*\**p*<0.001, \*\*\*\**p*<0.0001, one-way analysis of variance.

## Conclusion

A novel PSMA-guided PROTAC strategy was developed for the targeted degradation of POIs in prostate cancer cells with high PSMA expression. In particular, we designed a PSMA-guided AR degrader (**PSMA-ARD-203**) having an ester bond that could be hydrolyzed by intracellular hydrolases to release **ARD-203** for the degradation of AR. We confirmed that **PSMA-ARD-203** effectively degraded AR through the ubiquitin-proteasome pathway in a PSMA-dependent manner in PSMA-overexpressing prostate cancer cells. **PSMA-ARD-203** exhibited tumor tissue enrichment, leading to enhanced protein degradation in tumor tissues. To assess the generality of our guided strategy, we developed a PSMA-guided BET protein degrader (**PSMA-ARV-771**), which specially degraded BRD4 protein in prostate cancer cells having high PSMA expression. This study provides a generalizable platform for achieving the targeted delivery of PROTACs to effectively treat prostate cancer, thereby minimizing potential toxicity and side effects to normal tissues/cells while enhancing the *in vivo* antitumor efficacy.

## Materials and Methods

### Stability Assay of PSMA-ARD-203

The stability assay of **PSMA-ARD-203** (4 mg/mL) was performed using HPLC after incubating in PBS, cell culture media (DMEM) alone or DMEM plus 10% FBS at 37 °C for 2, 4, 8, 16 or 24 h. To precipitate proteins in FBS, the mixture was diluted with double volume of acetonitrile and centrifuged, followed by HPLC analysis for the top clear solution.

### Cell Lines and Cell Culture

LNCaP cells and 22Rv1 cells (Human prostate cancer cells) were purchased from Procell Life Science & Technology Co., Ltd. and cultured in RPMI1640 (Procell, PM150110) with 10% fetal bovine serum (Gibco, 51985-034) and 1% penicillin streptomycin. Vertebral Cancer of the Prostate (VCaP) cells, Human breast carcinoma (MCF-7) cells and Human embryonic kidney 293T (HEK-293T) cells were purchased from ATCC and maintained in Dulbecco’s modified Eagle’s medium (DMEM) containing 10% FBS and 1% penicillin streptomycin. MDA-MB-231 cells were obtained from Kunming Institute of Botany (Chinese Academy of Sciences) and maintained in DMEM/F12 containing 10% FBS and 1% penicillin streptomycin. J82 cells (Human transitional cell carcinoma of the bladder) and DU145 cells (Human prostate cancer cells) were cultured in MEM (NEAA) media (Procell, PM150410) containing 10% FBS and 1% penicillin streptomycin. J82 cells was obtained from Yantai Yuhuangding Hospital. DU145 cells were purchased from Procell Life Science & Technology Co., Ltd. Unless otherwise specifed, all cell cultures were grown at 5% CO_2_, 37 °C.

### Western Blot Assay

Whole cell lysates were collected using RIPA lysis buffer (Beyotime Biotechnology, Jiangsu, China) containing 1% protease inhibitor (Roche, 11697498001) and 1% phosphatase inhibitor (Roche, 4906837001). Lysates were centrifuged at 12,000 rpm at 4 °C for 10 min and the supernatant fraction was retained. Protein concentrations were quantified with BCA Protein Assay Kit (Thermo Fisher Scientific, A53226). Protein concentrations were quantified by BCA analysis, and equal amounts of protein were electrophoresed by 10% SDS-PAGE and transferred to polyvinylidene difluoride transfer membranes (PVDF, 0.45 μm) and incubated against different target antibodies at 4 °C overnight. Primary antibodies included β-Actin antibody (Abcam, ab20272), α-Tubulin antibody (Abcam, ab40742), PSMA antibody (Cell Signaling Technology, 12815S), AR antibody (Abcam, ab194196), membranes were subsequently washed in TBST and secondary antibodies conjugated to HRP were added in 5% milk and incubated for a minimum of 1 h at room temperature before developing with the ECL kit (Thermo Fisher Scientific, 34578).

### Transfection of PCa cell lines

LNCaP cells were transfected in Opti-MEM media (Gibco, 51985-034) using 50 nmol/L of each siRNA, and RNAi MAX transfection reagent (Invitrogen, 56532) according to manufacturer’s instructions. After 48 h, the cells were allowed to replace media and incubated with compounds for 12 h or 24 h. PSMA siRNAs were purchased from Sangon Biotech.

### Pharmacokinetic studies in Mice

Male BALB/c nude mice (4-6 weeks old) were implanted subcutaneously with 5 × 10^6^ 22Rv1 cells in Matrigel (ABW, 082724). When the tumor volume reached to about 300 mm^3^, the mice were randomized into groups and injected intravenously with the same concentration of drug (0.3 mmol/L). Blood samples were collected from the cheek at different time points after drug administration. Blood was collected using sodium heparin anticoagulation tubes and placed on wet ice for more than 15 min and centrifuged at 6000 r/min for 10 min, and the plasma was separated to collect the supernatant. Collect each animal tumor at the same time. The concentration of the sample was determined at each time points for each animal using LC–MS/MS. The follow-up test and analysis experiment were carried out by by Hefei Zhongke Precedo Biomedical Technology Co., Ltd. All experiments were conducted under a protocol approved by the Committee on the Ethics of Animal Experiments of Ocean University of China.

### *In vivo* therapeutic efficacy

Male BALB/c nude mice (5-6 weeks old) were purchased from SPF (Beijing) Biotechnology Co., Ltd. The mice were housed and maintained under SPF condition. Mice were implanted subcutaneously with 1×10^7^ LNCaP cells in Matrigel (ABW, 082724). When the tumor volume reached to 300 mm^3^, the mice were randomized into groups and injected intravenously with the same concentration of drugs (1 mmol/L) for three consecutive days. According to the experimental setting, the mice were sacrificed after 24 h, 48 h, and 72 h after the last dose, and tumor tissues were harvested for further analysis. All experiments were conducted under a protocol approved by the Committee on the Ethics of Animal Experiments of Ocean University of China.

### Cell Proliferation Assay

VCaP cells, 22Rv1 cells and LNCaP cells (5,000 cells per well) were dosed with compounds serially diluted 1:5 or 1:10 ranging from 1 μM for an 8-point dose curve for 72 h. Cell Titer-Glo Luminescent Cell Viability Assay (Beyotime Biotechnology, C0065L) was added, and the plate was read on a luminometer. Data were analyzed and plotted using GraphPad Prism software.

---

**Figure S1.**
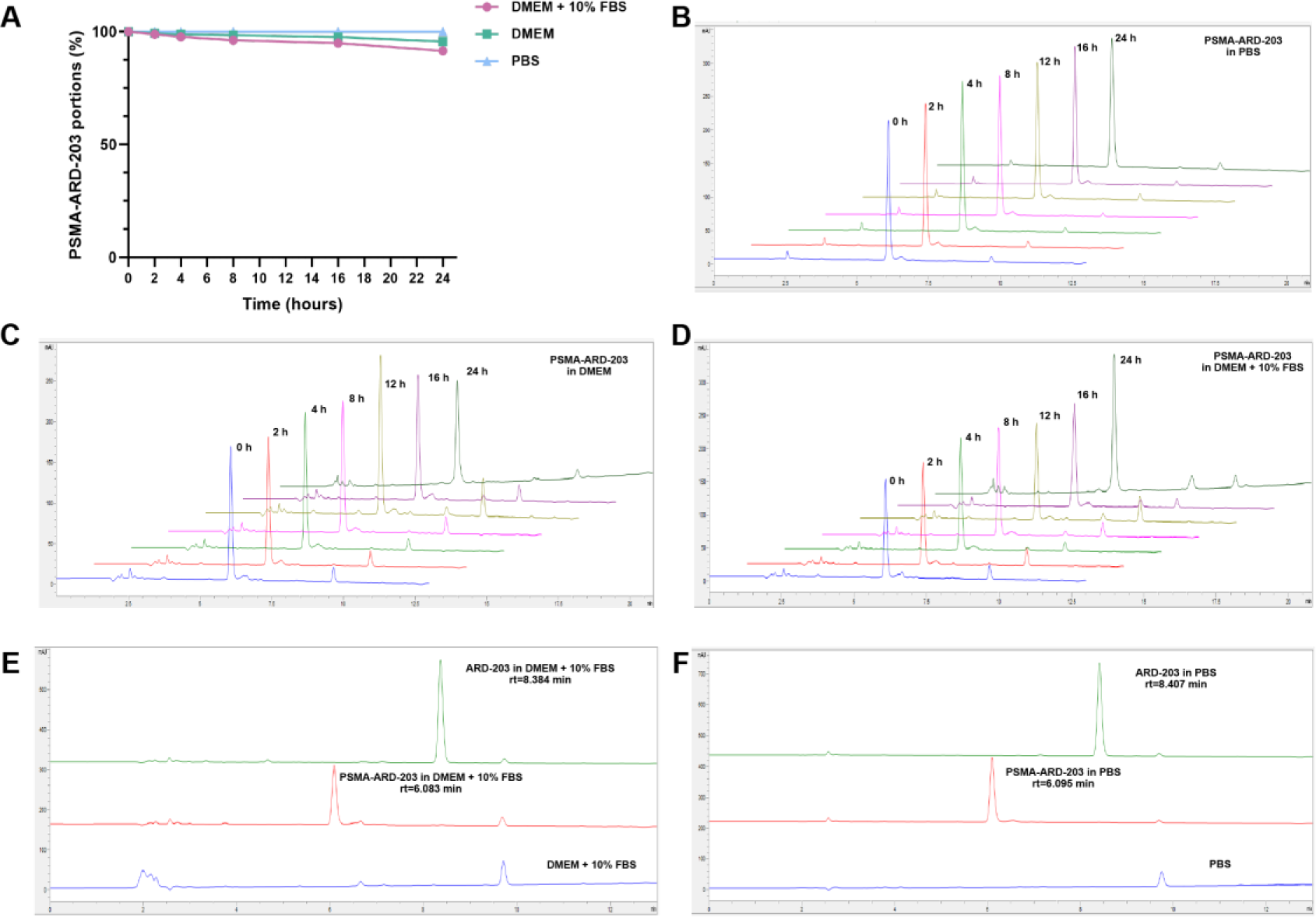
PSMA-ARD-203 is relatively stable in physiological conditions. (A-F) Stability assay of PSMA-ARD-203 in PBS, DMEM or DMEM+10% FBS with HPLC spectra.

**Figure S2.**
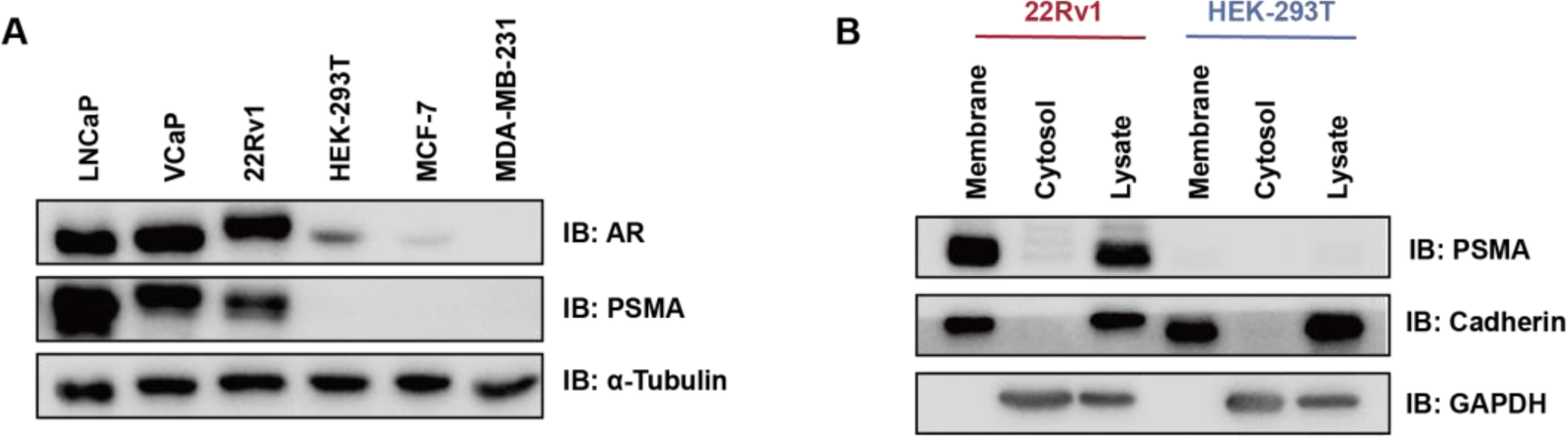
The expression of PSMA in different cell lines. (A) Western blot analysis of AR protein and PSMA levels in prostate cells or non-prostate cells. (B) Western blot analysis of cytosolic and membrane fractions from 22Rv1 cells and HEK-293T cells.

**Figure S3.**
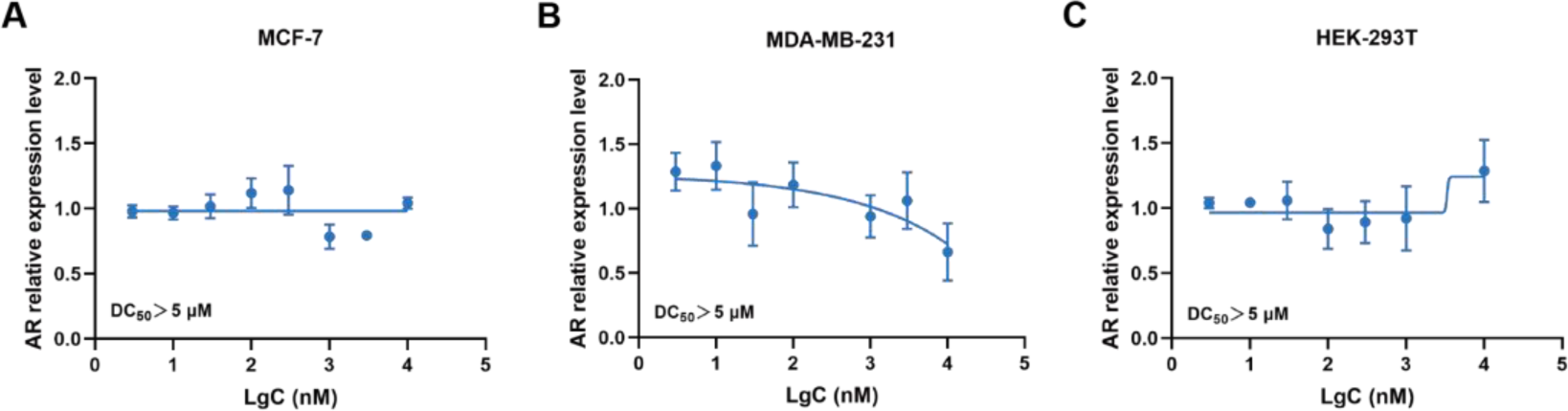
DC_50_ values of PSMA-ARD-203 based on quantification of AR protein levels normalized to α-Tubulin levels in non-prostate cell lines (n=3).

**Figure S4.**
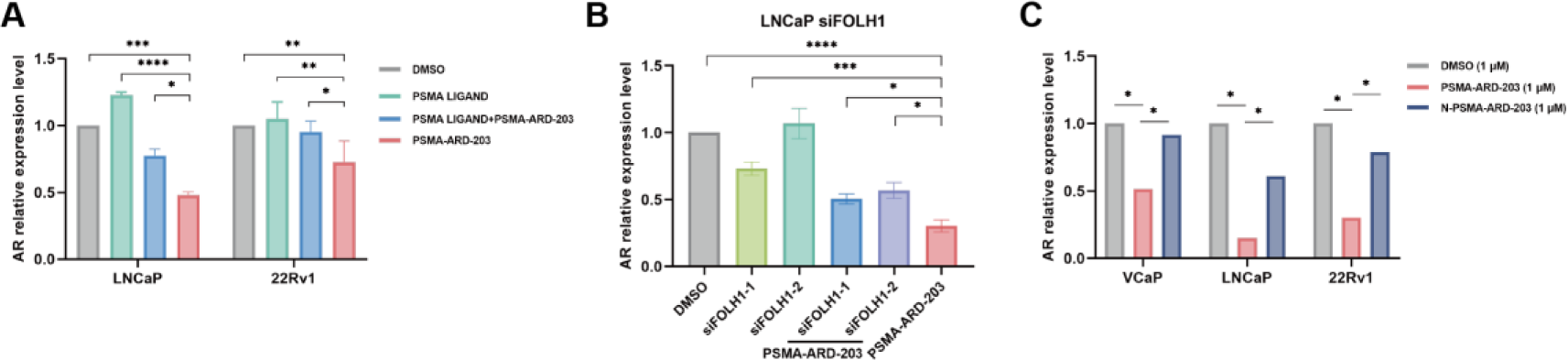
Quantification of AR protein levels of the Western blots in Figure 5 (n=3). Data are presented as mean ± standard error of the mean (n=3). \**p* <0.05, \*\**p*<0.01, \*\*\**p* <0.001, \*\*\*\**p*<0.0001, two-way ANOVA test.

**Figure S5.**
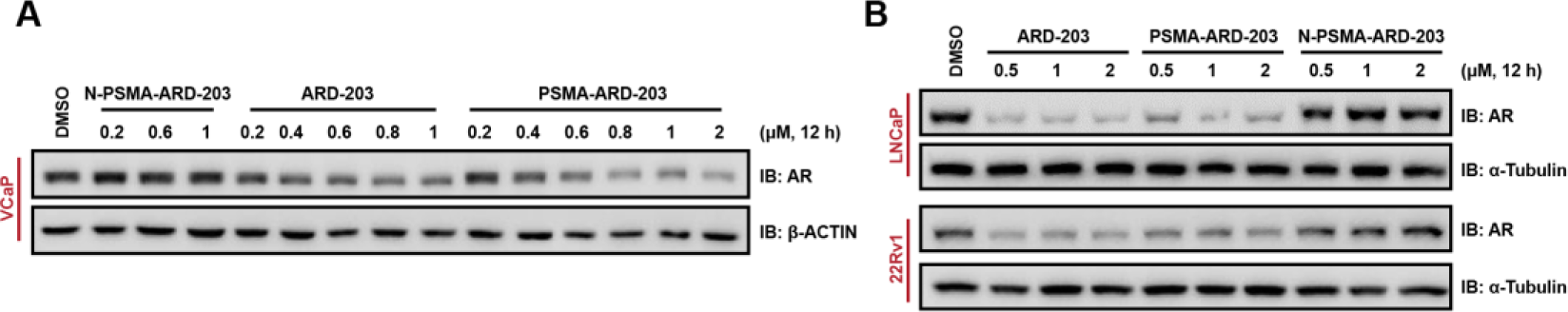
PSMA-ARD-203 degrades AR in PSMA-dependent manners. (A) Western blot analysis of AR in VCaP cells treated with indicated concentrations of ARD-203, PSMA-ARD-203 or N-PSMA-ARD-203 for 12 h. (B) Western blot analysis of AR in LNCaP cells and 22Rv1 cells treated with indicated concentrations of ARD-203, PSMA-ARD-203 or N-PSMA-ARD-203 for 12 h.

**Table S1.** PKs of PSMA-ARD-203 and ARD-203 in 22Rv1 Xenograft Tumors.

| Compound | Route | $T_{1/2}$ (h) | $T_{max}$ (h) | $C_{max}$ (ng/g) | $AUC_{0-t}$ (h·ng/g) | $MRT_{0-t}$ (h) | Cl (g/kg/h) |
| --- | --- | --- | --- | --- | --- | --- | --- |
| ARD-203 | I.V. | 13.11 ± 4.13 | 1.00 ± 0.00 | 179.00 ± 17.50 | 1289.00 ± 197.00 | 8.08 ± 3.08 | 2020.00 ± 382.90 |
| PSMA-ARD-203 | I.V. | 15.10 ± 4.39 | 1.00 ± 0.00 | 597.00 ± 332.00 | 7112.00 ± 1572.00 | 14.90 ± 6.20 | 628.00 ± 133.80 |
| Released-ARD-203 by PSMA-ARD-203 | I.V. | 24.80 ± 17.63 | 10.33 ± 12.10 | 100.00 ± 5.30 | 3223.00 ± 1182.00 | 24.18 ± 8.35 | 1330.00 ± 652.80 |

<sup>a</sup> $C_{max}$ , maximum drug concentration; $AUC_{0-t}$ , area-under-the-curve between 0 and 96 h; Cl = plasma clearance rate; $T_{1/2}$ = terminal half-life, $T_{max}$ = the time take to reach $C_{max}$ ; $MRT_{0-t}$ , mean residence time; I.V., intravenous administration. Data are shown as the mean ± standard deviation (n=3).

**Figure S6.**
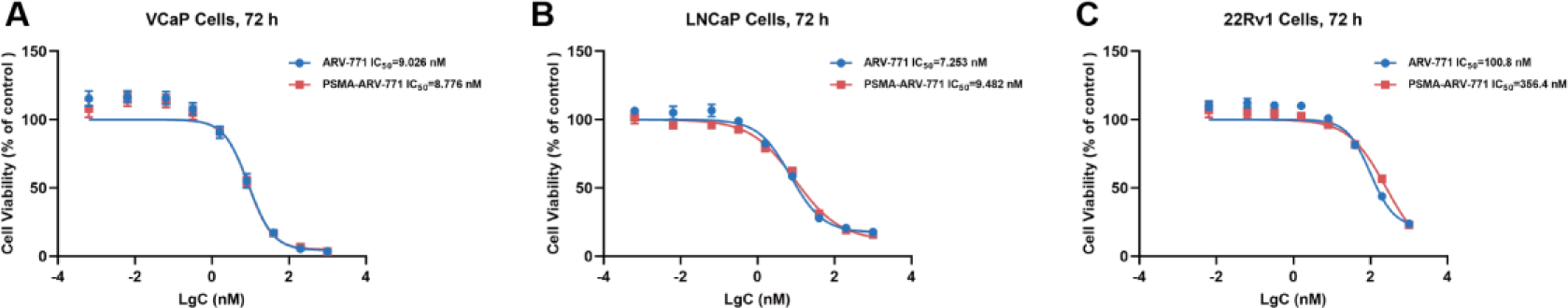
Cell viability of VCaP, LNCaP and 22Rv1 cells after treatment with PSMA-ARV-771 or ARV-771 for 72 h.

